# Long-read transcriptome sequencing analysis with IsoTools

**DOI:** 10.1101/2021.07.13.452091

**Authors:** Matthias Lienhard, Twan van den Beucken, Bernd Timmermann, Myriam Hochradel, Stefan Boerno, Florian Caiment, Martin Vingron, Ralf Herwig

**Author notes:** To whom correspondence should be addressed., ML, RH.

## Abstract

Long-read transcriptome sequencing (LRTS) holds the promise to boost our understanding of alternative splicing. Recent advances in accuracy and throughput have diminished the major limitations and enabled the direct quantification of isoforms. Considering the complexity of the data and the broad range of potential applications, it is clear that highly flexible, accurate analysis tools are crucial. Here, we present IsoTools, a comprehensive Python-based analysis package, for the improvement of alternative and differential splicing analysis. IsoTools provides a comprehensive data structure that integrates genomic information from LRTS transcripts together with the reference annotation, and enables broad functionality to quality control, visualize and analyze the data. Additionally, we implemented a graph-based method for the identification of alternative splicing events and a statistical approach based on the beta binomial distribution for the detection of differential events. To demonstrate our methods, we generated PacBio Iso-Seq data of human hepatocytes treated with the HDAC inhibitor valproic acid, a compound known to induce widespread transcriptional changes. Contrasted with short read RNA-Seq of the same samples, this analysis shows that LRTS provides valuable additional insights for a better understanding of alternative splicing, in particular with respect to complex novel and differential splicing events. IsoTools is made available for the community along with extensive documentation at https://github.com/MatthiasLienhard/isotools.

## Introduction

Long Read Transcriptome Sequencing (LRTS) allows for full-length sequencing of expressed transcripts. In contrast to short-read RNA-Seq, this technology does not require fragmentation of transcripts and thereby avoids introduction of errors due to alignment ambiguity and other technical artefacts. This is particularly relevant for the analysis of complex splicing events since short reads cannot be assigned reliably to transcript isoforms over longer genomic distances [1, 2, 3].

With the LRTS Iso-Seq protocol from PacBio, besides direct cDNA/RNA sequencing by Oxford Nanopore, a technology has emerged that holds the promise to improve isoform identification and quantification. While hybrid sequencing approaches have been suggested in the past to combine detection (LRTS) and quantification (RNA-seq) of isoforms [4], recent advances in accuracy and throughput facilitate direct quantification of isoforms and splicing events from LRTS data alone.

LRTS provides a broad range of potential use cases for both model and non-model organisms. For poorly annotated non-model organisms the technology has been applied to facilitate the identification of relevant gene structures and coding regions [5, 6]. Common use cases for human samples and other model organisms include the discovery and characterization of novel genes, transcripts and alternative splicing events, as well as facilitating the quantification of isoform expression, either with or without integrating short-read RNA-Seq data [7, 8]. Very recently, LRTS has been combined with single-cell technology, to explore and characterize splicing on the level of cell types [9, 10, 11].

Recent studies suggested the possibility of direct quantitative interpretation of LRTS data on transcript isoform level, bypassing the error prone iterative assignment of short reads [12, 13, 14]. On gene level, a fair correlation of 0.65 and 0.8 between long read IsoSeq and short read RNA-Seq quantification was found for human and mouse cortex samples. However, on isoform level this correlation dropped to 0.36 and 0.41 [13].

Besides the quantification of isoforms, differential splicing can be analysed on the level of alternative splicing events (ASEs). The most common definition of these ASEs are based on bubble structures in splice graphs [15], usually constructed from annotated gene models. Differential analysis of ASEs is common with short-read RNA-Seq [16, 17], and offers advantages due to reduced complexity and improved interpretability. However, as short reads do not cover the complete events, particularly for complex events that are connected over longer genomic distances, this approach also suffers from ambiguity.

For many of the common use cases, specific tools have been provided. Most prominently, Functional IsoTranscriptomics (FIT) is a comprehensive framework, including SQANTI3 [18], IsoAnnot and TappAS for quality control, isoform and splice junction annotation and classification, and prediction of functional implications of alternative isoforms. These tools provide convenient standardized analysis pipelines, facilitating characterization and functional interpretation of alternative splicing by open reading frame (ORF) prediction and differential isoform expression analysis between samples by integrating RNA-seq data for isoform quantification. As an alternative, TAMA focuses on filtering sequencing noise and detecting novel expressed transcripts, in particular lncRNAs [19]. TALON was developed as the ENCODE4 pipeline for long-read sequencing, and provides similar functionality to SQANTI3 [12].The Swan library [14], which is part of the ENCODE4 TALON pipeline, implements a statistical test based on a negative binomial model for differential expression analysis on isoform level using LRTS.

Considering the complexity of the data and the broad spectrum of potential use cases, it is clear that standardized analysis tools may become limiting. To unravel the full potential of LRTS, researchers need to be able to fully explore the data at all scales, ranging from single nucleotide information over transcript and gene level, to transcriptome-wide statistics. With this in mind, we developed IsoTools, a flexible Python framework for the analysis of PacBio Iso-Seq data, which implements data structures integrating all relevant information from LRTS transcripts and reference annotation, together with broad analysis functionality to explore, analyze, and interpret the data.

In addition to this framework, we provide tutorials and Jupyter notebooks covering the most relevant use cases, including analysis of differential splicing events between samples solely based on long-read sequencing or in combination with short-read sequencing. The documentation also includes the complete API reference, facilitating custom analysis approaches. IsoTools was extensively tested on published LRTS data from PacBio Sequel I and Sequel II. To further demonstrate the utility of the workflows, we generated LRTS data of human primary hepatocytes, treated with the HDAC inhibitor valproic acid (VPA) with latest technology (PacBio Sequel II Iso-Seq). VPA is known to induce a wide spectrum of changes in the epigenome and the transcriptome leading to differential isoform usage [20]. Identified novel and differential splicing events from several categories including exon skipping, mutually exclusive exons, and novel poly-A sites were validated with (short-read) RNA-Seq on the same samples. We further investigated the *in-vivo* relevance of detected alternative splicing by confirming expression in healthy human liver tissue.

## Results

### IsoTools provides full control over all stages of LRTS data analysis

One challenge of LRTS analysis is posed by the wide range of information on different scales, ranging from single nucleotides (e.g. variant information) over exon information, alternative splicing events, transcript isoforms, genes, to transcriptome-wide statistics, e.g. on transcript length distribution. We address this challenge by implementing an efficient tree-based data structure, facilitating access to all relevant information by genomic positions as well as by names or identifiers. In order to provide the user full control over the data, IsoTools can import these information from aligned long read files and reference annotation. During import of the long reads, the transcripts are assigned to reference genes and categorized. After import, the complete gene models are encoded as segment graphs [21], where nodes represent disjoint exonic segments, while edges imply that the two exonic segments succeed one another within one of the transcripts (see Methods). This segment graph is an efficient way to store complex splicing structure of a gene and facilitates a range of algorithms to characterize and compare transcripts, as well as to define alternative splicing events. Figure 1 provides an overview on the internal data structure of the IsoTools framework.

**Figure 1:**
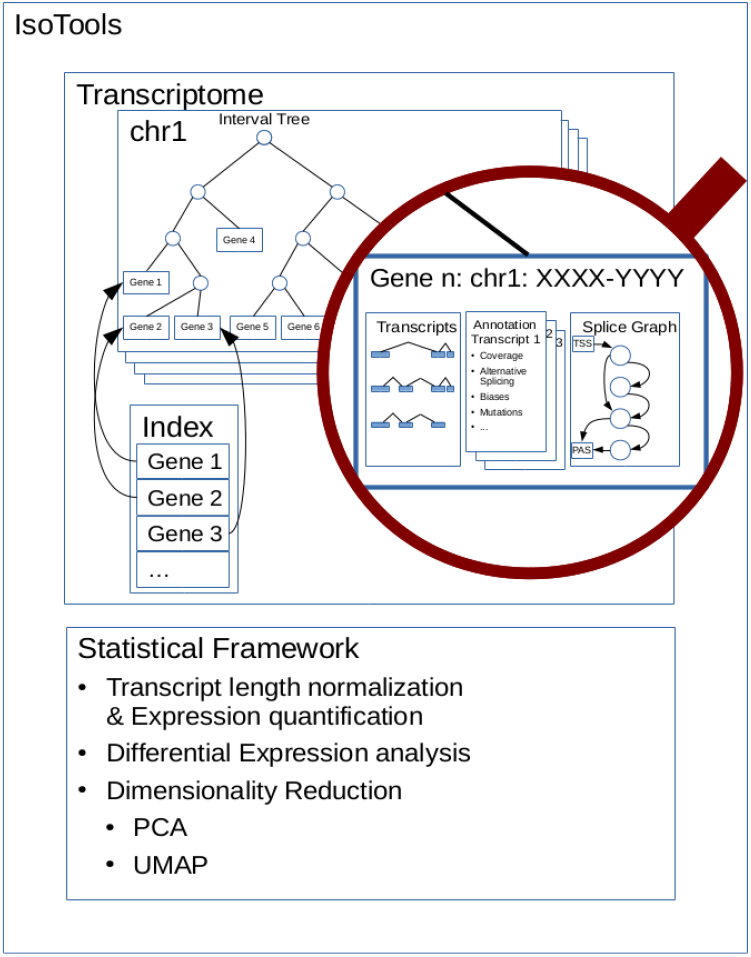
The internal data structure of the IsoTools framework

To demonstrate a basic LRTS workflow using IsoTools, we produced a PacBio IsoSeq dataset from human hepatocytes treated with the HDAC class I and II inhibitor valproic acid (VPA) and controls (CTL), yielding 2,615,181 and 4,200,885 aligned full-length non chimeric poly-A reads, respectively.

### Saturation analysis estimates required sequencing depth

Prior to LRTS data generation it is of general interest to assess the required depth of sequencing for recovering transcripts of interest. We developed a general saturation model based on 3 relevant parameters. First, the required sequencing depth depends on the cellular concentration of the transcript in the samples. A highly expressed transcript is more likely to be covered compared to a transcript with few RNA molecules per cell. Second, the minimum required coverage to confidently call a transcript impacts the chance of a transcript to be discovered. The PacBio IsoSeq clustering pipeline requires two copies of a transcript to report it. However, depending on the application, it may be appropriate to reduce or increase this threshold. As third factor, transcript discovery depends on the sequencing depth. Assuming sufficient library complexity, the more transcripts are sequenced, the higher is the chance of sequencing one particular transcripts.

Neglecting potential biases and assuming linear relation between RNA concentration and sampling probability, the probability of observing a transcript can be modeled by the negative binomial distribution. Figure 2 provides the probability of observing transcripts at different expression levels and at different detection thresholds, depending on the read coverage. According to this model, the probability of observing a transcript expressed at one TPM with at least two reads is 93% for the CTL sample and 74.7% for the VPA treated sample (intersection of orange line and vertical lines in Figure 2B). For transcripts at a cellular concentration of 2 TPM, the probability of observing at least two reads is close to 100% for both samples, indicating saturation of transcript discovery at this concentration and detection threshold.

**Figure 2:**
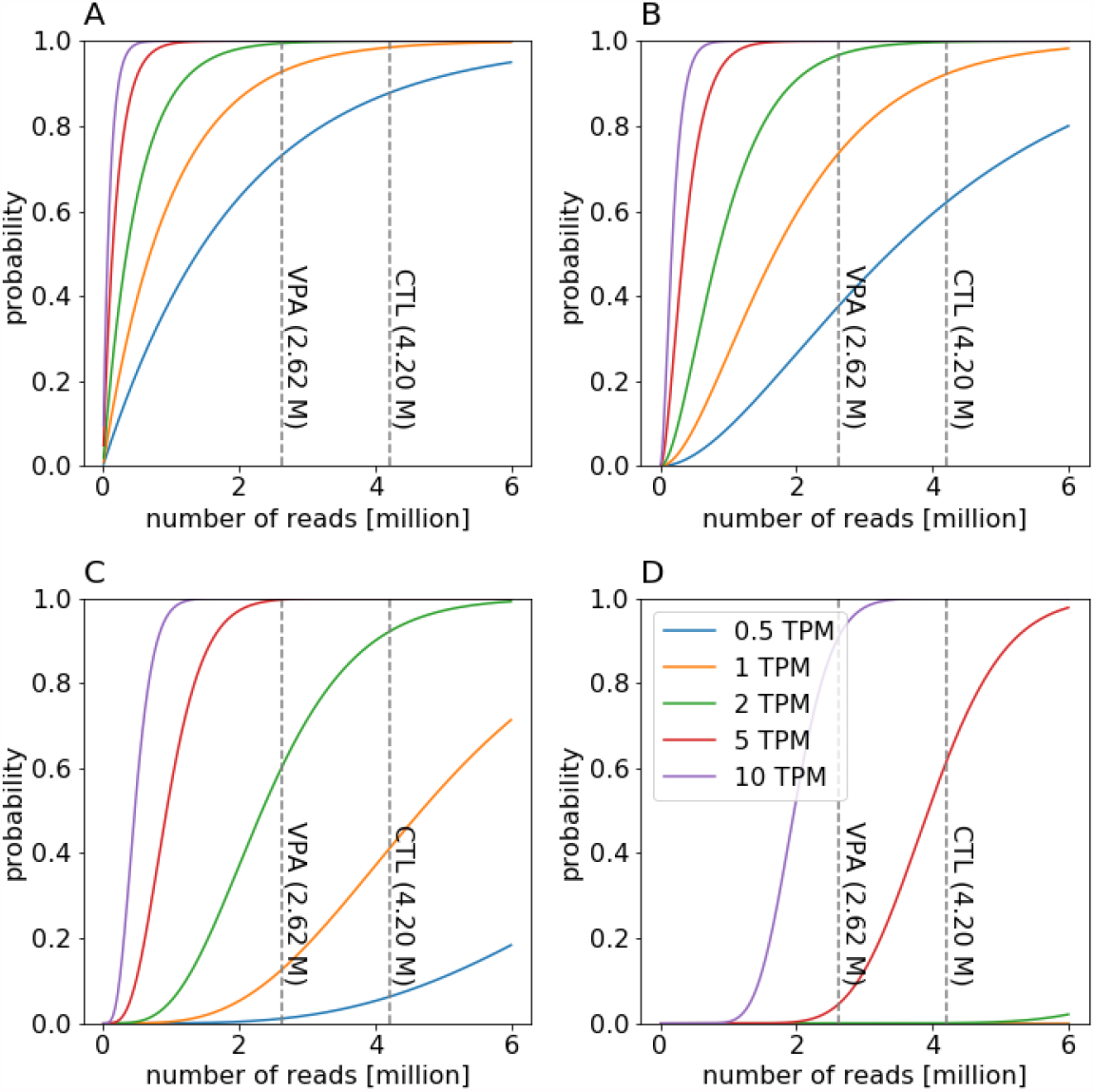
Probability of observing a transcript depending on the cellular concentration from 0.5 (blue line) to 10 TPM (purple line) and sequencing depth at detection threshold of 1 read (A), 2 reads (B), 5 reads (C) and 20 reads (D). Dashed horizontal lines represent the sequencing depth of the VPA and CTL IsoSeq samples.

### Quality control measures allow filtering for technical artifacts

Although LRTS holds the promise to boost alternative splicing, several technical artefacts have been identified that might bias or impair LRTS analysis. In order to control for these artefacts and to filter related transcripts in a flexible and efficient way, IsoTools provides different quality control (QC) measures (Figure 3).

**Figure 3:**
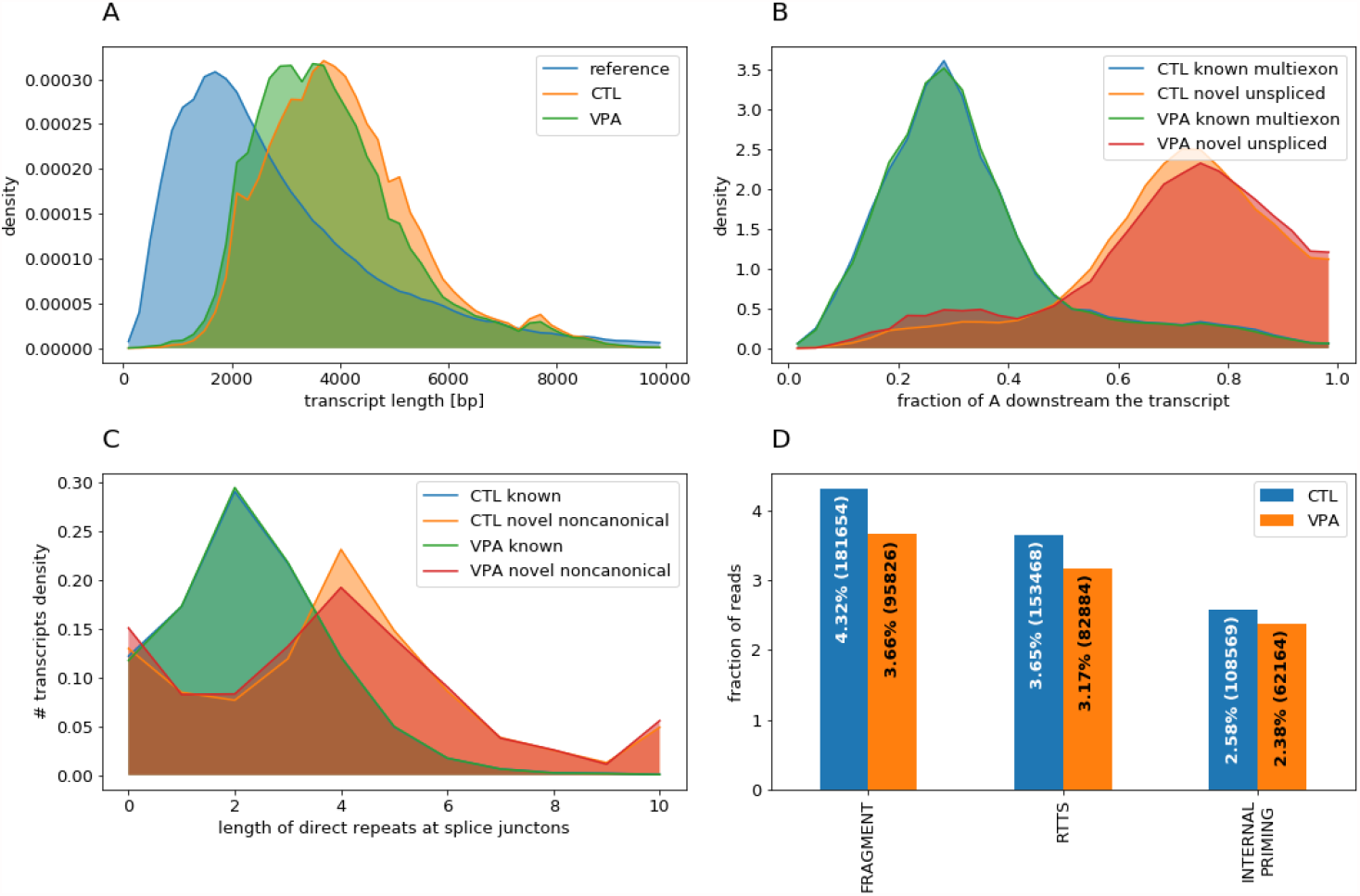
Quality control metrics. A) Read length distribution compared to level 1 GENCODE transcripts. B) A content downstream of novel unspliced transcripts and reference matching multi-exon transcripts. C) Direct repeat length at intron boundaries of gencode transcripts and novel non-canonical splice junctions. D) Fraction of reads affected by one of the three artifacts.

#### The transcript length distribution

reflects the depletion of transcripts depending on their length. It should be comparable between samples and can be compared to the length distribution of all transcripts in the reference annotation. Quite surprisingly, we noticed that the transcripts sequenced with Sequel2 are longer compared to the annotated transcripts, suggesting a depletion of shorter transcripts. (Figure 3A)

#### Technical artifacts

are indicated with different QC measures. Here, we consider three kinds of technical artefacts known to occur in LRTS data [18]:

- Genomic stretches of adenosines may be bound by the poly-A primer, inducing sequencing of genomic templates, which are not spliced and thus yield a single exon transcript in LRTS. This effect is called **internal priming**.
- The ability of the reverse transcriptase to switch between templates (**RTTS**) is exploited to anneal the primers during the SMRT sequencing. However, this switching can also occur unintended within or between templates, resulting in incomplete or chimeric templates. This template switching preferably occurs at short direct repeats.
- Transcripts may be **truncated** during library preparation, resulting in transcript fragments. Due to poly-A priming, 3’ fragments are unlikely to be sequenced, but 5’ fragments yield incomplete transcripts with apparent novel transcription start sites.

To control for these artefacts, IsoTools facilitates the definition of filters to tag affected transcripts. For subsequent analyses, visualizations, and data exports, the user can filter the transcripts according to these tags. In order to find reasonable thresholds for the filter definitions, we compared corresponding metrics from the most credible GENCODE transcript with support level 1 to the most suspicious transcripts identified by LRTS. However, all filter expressions may be adapted or extended by the user.

To identify internal priming, we monitor the fraction of adenosines in the genomic sequence 30 bases downstream of the transcript. For the highly credible (support level 1) GENCODE transcripts we find a downstream adenosine content distributed with a single mode around 25% adenosines. However, when looking at single exon genes that do not overlap any reference genes, we observed a second mode at 70% adenosines. For the CTL and VPA samples, 55.5% and 48.3% of all “novel” single-exon transcripts feature > 50% adenosines respectively (Figure 3B). This suggests, that 50% downstream adenosienes is a reasonable threshold to mark internal priming. To select for putative RTTS sites, we screen for introns without reference support where both donor and acceptor sites are within reference exons, and which do not feature a canonical splice site. At the boundaries of these introns, we compute the length of direct exact repeats and compare it to the repeat length at regular introns of the high confidence GENCODE transcripts (3C). While the putative RTTS sites feature a slightly longer average repeat length of 5 bases compared to 2 for the regular introns, this difference does not allow to define a threshold that would separate the majority of putative RTTS transcripts from the credible transcripts. Therefore we simply label transcripts as RTTS if they have a non-canonical, non-reference intron, e.g. both donor and acceptor are not annotated and the sequence at the splice site is not ‘GT-AG’.

Without additional experimental evidence, there is no way of differentiating true alternative start sites from potential truncations. All transcripts that start or end within an internal exon of another transcript, but share all other splice sites with this transcript, may be considered potential fragments.

According to the definitions above, 10.5 % and 9.2 % of the reads are affected by technical artefacts in the CTL and VPA hepatocytes samples. However, the largest fraction is contributed by potential transcript fragments, affecting 4.3% and 3.7% of the reads, for CTL and VPA. This is followed by RTTS (3.7% and 3.2%) and internal priming (2.6% and 2.4%) (Figure 3D).

In order to assess potential truncations, we took additional evidence from publicly available CAGE data into consideration. To this end, we downloaded CAGE TSS peaks of HepG2 cells from ENCODE [22]. 9.9% of the potential fragments feature overlapped CAGE peaks in HepG2 cells from ENCODE, compared to 76.9% of all transcripts expressed with at least 2 reads that correspond to high confidence GENCODE transcripts, suggesting that many of the fragments are indeed truncated transcripts. However, the remaining 9.9% are good candidates for novel transcription start sites, and might therefore be of particular interest and should not be discarded. Also, the lack of CAGE peaks is not sufficient evidence to rule out a TSS. Furthermore the ENCODE cell line samples are quite different from the primary hepatocytes used in this study.

In order to cope with such ambiguity, IsoTools does not filter out potential technical artefacts, but rather tags these cases and lets the user decide in which situations transcripts associated with biases are considered or neglected.

### Chimeric alignments identify fusion transcripts

An interesting aspect in analyzing LRTS data are chimeric alignments. These alignments can have several potential causes, of both technical and biological origins and IsoTools provides methods to distinguish these cases.

First, large introns > 200 kb often can not be mapped by the alignment tool and are reported as chimeric alignments. Also, RTTS between templates yields chimeric cDNA fragments. Further, during PCR amplification, templates may get joined together through transcriptional slippage [23], yielding apparently fused transcripts. However, there are also interesting biological effects resulting in chimeric alignments: mRNAs can be spliced back, yielding circular RNA molecules [24]. Since circular RNAs do not contain poly-A tails, these cases are usually not covered by LRTS and only appear in specific situations, such as due to internal priming. Further, fusion transcripts result in chimeric alignments. Fusion transcripts may either emerge from genomic rearrangements or trans splicing, and are commonly found in cancer cells [25]. In principle, LRTS provides great means to not only identifying these fusion transcripts but also revealing the fraction of fused vs. unfused transcripts as well as the transcript isoforms involved.

Chimeric alignments mapping on the same strand and less than 1 megabase apart can be considered as large introns, and hence the parts can be chained to a single alignment. To separate the other cases is challenging and we should consider additional evidence of the fusions and critically evaluate the plausibility. To facilitate these considerations, IsoTools extracts the breakpoints, gene annotations and sequencing coverages of all additional chimeric alignments for further inspection as a table (Suppl. Table 1).

In total, we found 30,561 and 66,293 chimeric alignments for the VPA and CTL samples respectively, of which 3,610 and 4,688 could be chained to a consecutive alignment, and thus are likely caused by long introns. To reduce the impact of the technical artefacts, we focused on chimeric alignments that were supported by at least 2 reads, leaving 1,786 and 3,737 reads respectively. Of those, the majority (1,287 and 2,144 reads) have both nonconsecutive parts of the alignment at the MALAT1 lncRNA locus, but the second part of the alignment is upstream of the first part. Such alignments could be explained by backsplicing, however, as the respective junctions have no short-read support, they are more likely technical artifacts.

The remaining chimeric reads cover 21 breakpoints with at least 10 reads over both samples, involving 30 different genes. Despite the fair coverage, none of these chimeric alignments show significant short-read support, making RTTS and/or transcriptional slippage during the PCR step more likely explanations. This implies that potential fusion transcript candidates identified by IsoSeq require careful investigation and validation, also for samples where gene fusions might be more plausible.

### Refined classification scheme facilitates biological interpretation of novel transcript isoforms

In order to assess the nature of novel transcripts, Sqanti [18] introduced a classification scheme based on a comparison to the reference transcriptome. The authors distinguish full splice matches (FSM), incomplete splice matches (ISM) matching consecutive splice junctions, novel in catalog (NIC) containing exclusively annotated splice sites, and novel not in catalog (NNC) containing at least one novel splice site. We adopted this scheme, but extended it by additionally assigning specific subcategories to the transcripts that enable direct biological interpretation, and thus inference of underlying biological principles.

ISMs correspond to fragments of reference transcripts, and IsoTools distinguishes “5’ fragments”, “3’ fragments” and “mono-exons”. NIC transcripts are sub-classified in “exon skipping”, “intron retention”, “novel combinations” of known splice junctions and other “novel junctions”, e.g. both splice sites are annotated, but not used by the same junction according to the reference. Further, if the first or last exon shares its splice site with an internal reference exon, the transcript is classified as “novel exonic TSS” or “novel exonic PAS”. NNC contains transcripts with “novel 5’ splice sites” (splice donors), “novel 3’ splice sites” (splice acceptors), or “novel exons”, not overlapping any reference exon. If a novel exon is the first/last exon of the transcript, it is classified as “novel intronic TSS/PAS”. If a transcript includes splice sites from more than one reference gene it is classified as “readthrough fusion”. Finally, transcripts not overlapping any splice junctions from the reference are classified as novel genes. Here, we follow the subclassification of Sqanti and distinguish “genic genomic” (implying exonic overlap with a reference gene), “intronic”, “antisense” and “intergenic” transcripts.

Note, while the Sqanti categories are mutually exclusive, a transcript may be assigned to several subcategories. Suppl. Figure S1 provides an overview of the different categories. To find these subcategories we compare the exon structure of an LRTS transcript to the segment graph of the reference gene. This implementation facilitates the definition of a set of rules to identify the relevant subcategories.

In the primary hepatocyte LRTS, 72.5% and 73.5% of the reads (after filtering for technical artefacts) fully match known transcripts (FSM) while the most prevalent category for novel transcripts are novel combinations of known splice junctions, affecting 10.1% and 9.5% of the reads, for CTL and VPA treated samples respectively (Figure 4). While the high fraction of transcripts in this non-reference category may be surprising, the evidence is clear and convincing in this case and reflects the incompleteness of reference annotation. One of the highest expressed transcripts in this category is a novel isoform of the SPTBN1 gene, which combines the promoter of the isoform SPTBN1-201 with the poly-A site of SPTBN1-202 and SPTBN1-207, after sharing 30 intermediate exons with both transcript variants. (Suppl. Figure S2). The novel transcript is covered by 3306 and 1620 IsoSeq reads, about 30% of all reads of that gene. Note that both alternative splicing events affect the coding region, potentially yielding a novel protein sequence. As the next most frequent categories of novel transcripts, we observed novel exon skipping as well as 5’ and 3’ alternative splice sites. However, we found that a large part within this categories must be attributed to misalignment of short exons < 30 bases, which are aligned to either of the neighboring exon boundaries. This issue affects up to 20% of the transcripts in these categories, and more than 50% of the highly covered transcripts with more than 50 IsoSeq reads (Suppl. Figure S3).

**Figure 4:**
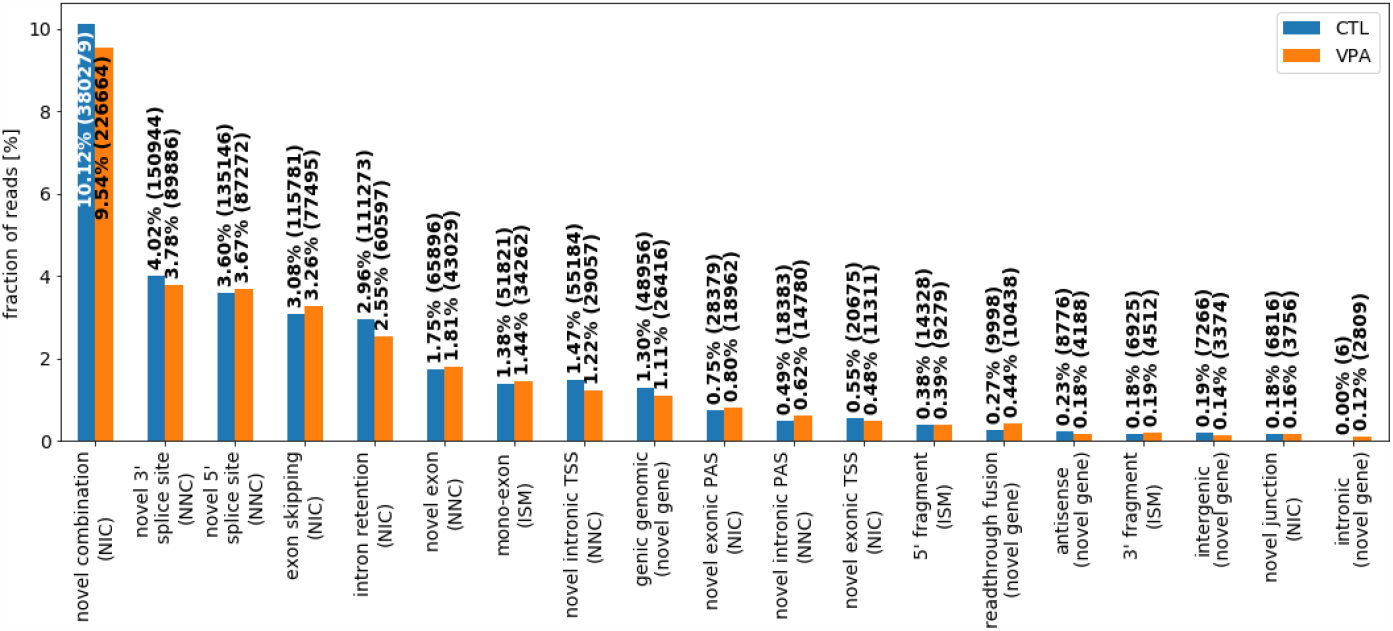
Alternative splicing classification of the IsoSeq reads not fully matching reference annotation for CTL and VPA treated hepatocyte samples.

One example of a novel skipping event which is likely not a technical artefact affects the 102 bp exon 7 of PATL1, which is skipped in 28.8% and 21.5% of the transcripts in CTL and VPA respectively (Suppl. Figure S4). PATL1 is involved in mRNA degeneration, and the skipped exon overlaps the protein domain involved in RNA-binding, according to UniProt [26]. Junction coverage of short-read data from the same samples confirm exon skipping of a similar proportion of transcripts. Notably, the exon skipping event can also be found in short read RNA-Seq of human liver samples, suggesting *in-vivo* relevance of the novel isoform.

3.0% and 2.6% of transcripts feature retained introns, which may be due to incomplete processing of pre-mRNA. The categories “5’ fragment”, “novel exonic TSS” and “novel intronic TSS” all describe potential novel alternative transcription start sites. We checked these sites for CAGE support from the HepG2 cells and found 11.7%, 10.1% and 31.1% of the transcript starts overlapping CAGE peaks for the 3 categories respectively, compared to 76.9% for the full splice match transcripts. While novel transcription starts overlapping exons can also be explained by fragmentation, novel intronic TSSs provide the best evidence for true novel TSS. All transcripts found covered by at least 10 reads in the primary hepatocyte samples, along with their novelty classification, are listed in Suppl. Table 2.

### LRTS provides quantitative information on expression level

Recent advances with the new Sequel II system allow direct quantitative interpretation of the transcript counts. Compared to the predecessor, the maximal CCS read length has increased, and thus the technology is capable of covering the majority of the transcriptome without depleting longer transcripts. Previously, this bias was countered with enrichment of long fragments during library preparation, disturbing the quantitative signal of transcript read counts. In addition, the throughput has increased 8-fold at comparable cost and time requirements, facilitating the required sequencing depth for quantitative interpretation of the read counts.

After basic normalization for sequencing depth, both hepatocyte samples had a similar distribution of read counts over transcripts (Suppl. Figure S5). Consistent with recent results [13], we also found good agreement between RNA-Seq gene expression levels and Sequel II IsoSeq coverage (*r* = 0.756 and 0.765 for CTL and VPA), underlining the quantification performance (Figure 5). On transcript isoform level, the correlation dropped considerably (*r* = 0.421 and 0.427), also confirming previous results [13]. A plausible explanation for the reduced accordance on transcript level is the uncertainty in assigning ambiguous short RNA-Seq reads, and the errors introduced due to violated assumptions of uniform read coverage [27] and imperfect reference model [28].

**Figure 5:**
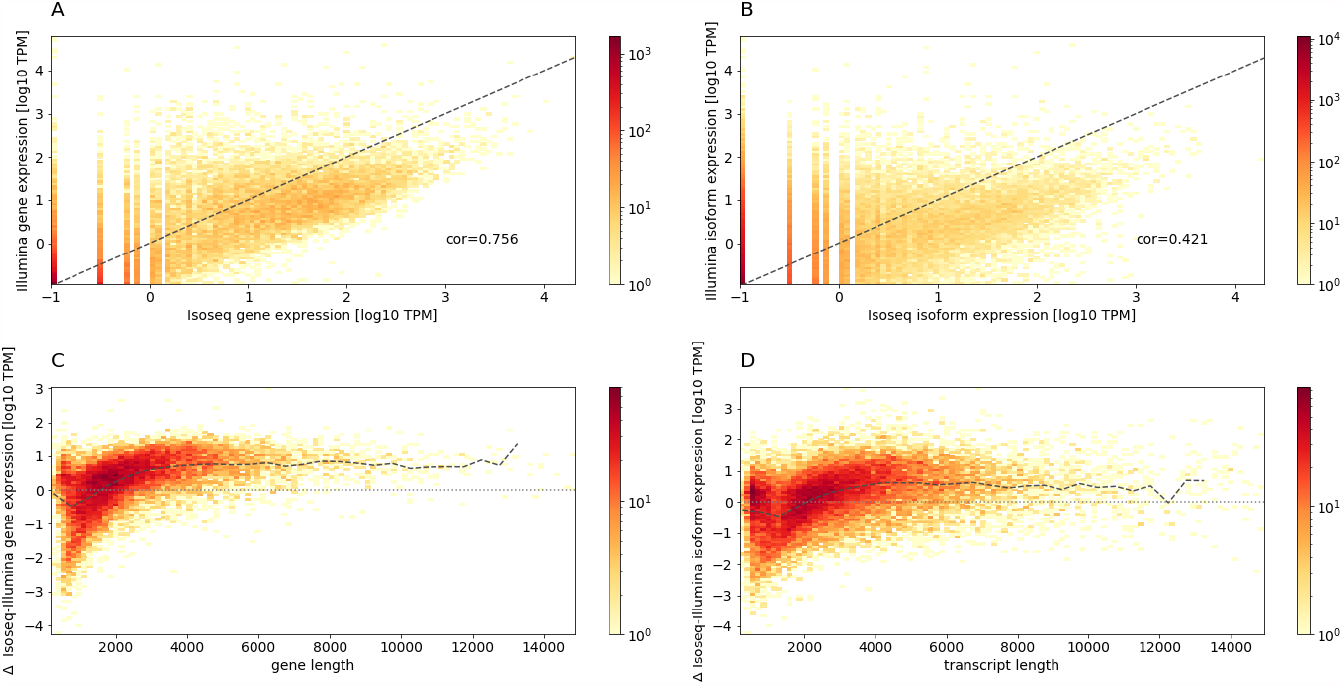
Comparison of expression levels derived from Illumina RNA-Seq and IsoSeq on gene level (A) and transcript isoform level (B). RNA-Seq vs IsoSeq expression log10 difference depending on transcript length on gene level (C) and transcript level (D). Dashed line represents the binned average.

To investigate a potential transcript length bias, we analysed the deviance of IsoSeq derived expression levels from RNA-Seq derived expression levels, depending on the transcript length. For transcripts > 3, 000 bases, this deviance was constant, reflecting good agreement between RNA-Seq and IsoSeq. However, for shorter transcripts, IsoSeq yields lower expression levels, suggesting a depletion of short transcripts within our IsoSeq data. This finding is in line with the comparison of transcript length to the reference annotation (Figure 3A).

### Alternative splicing events facilitate interpretation of complex splicing

Even though LRTS serves as a direct approach to transcript reconstruction and quantification, the statistical analysis and interpretation on this level is not trivial because of the large number of potential different transcripts. Similar to RNA-seq analysis it might thus be convenient to decompose the splicing landscape of a gene into alternative splicing events (ASEs).

To this end, we implemented a method to detect ASEs based on segment graphs (Figure 6 A and B). This approach allows to isolate the individual events that distinguish the isoforms, and hence break down the complexity. The relative expression of these events is quantified with the percent splice index (PSI), which is the fraction of reads supporting the alternative (see Methods). Our definition additionally subdivides and classifies splicing events according to the underlying molecular principles as exon skipping (ES), intron retention (IR), 5’ and 3’ alternative splicing (5AS and 3AS), and mutually exclusive exons (ME) (Figure 6 C).

**Figure 6:**
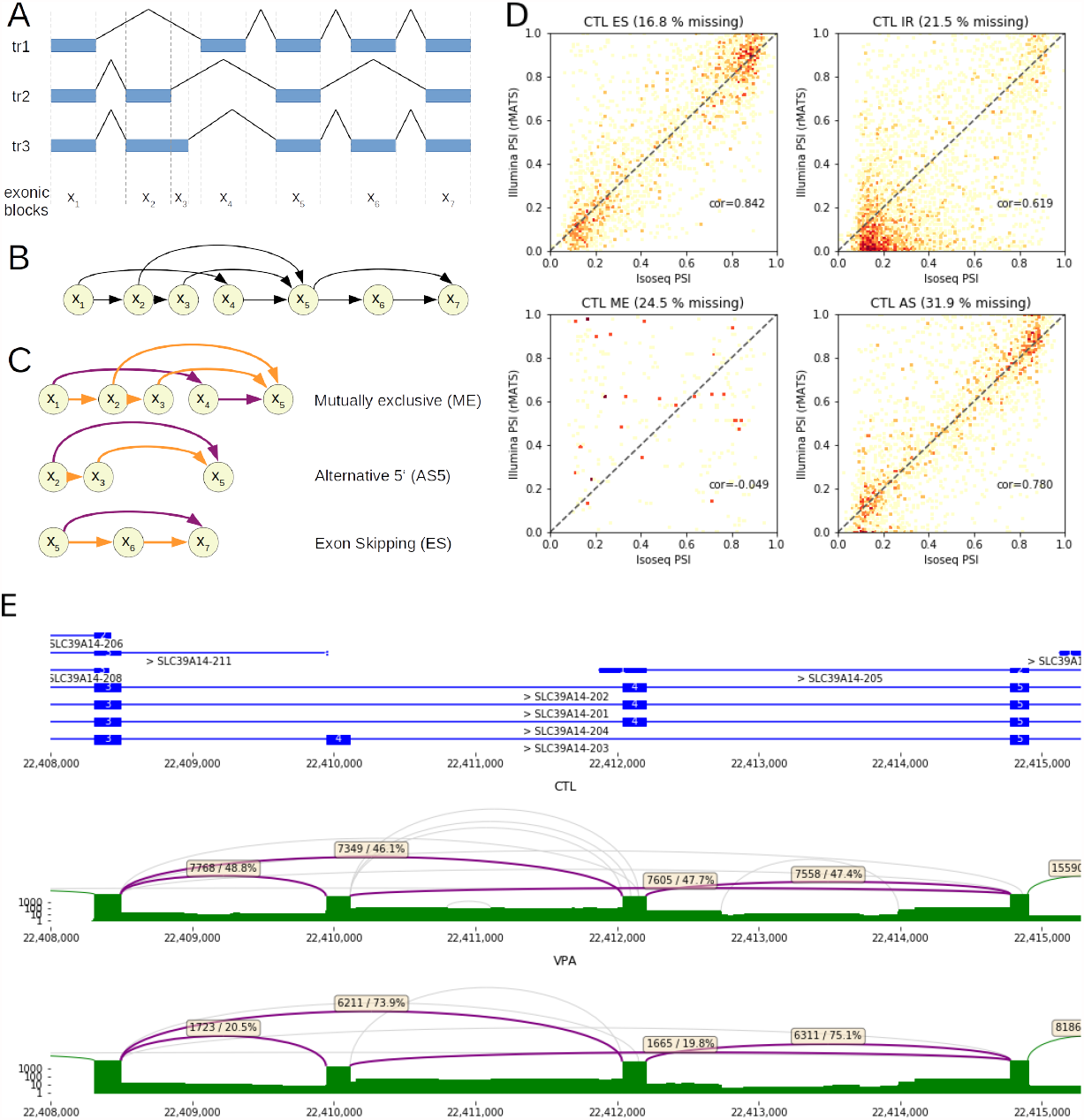
Splice Bubble decomposition of the Segment Graph to identify alternative splicing events (ASEs). A) Exemplary gene with 3 transcripts and B) the corresponding Segment Graph. C) The 3 ASEs and corresponding categories, defined by the Segment Graph. Note that the ME event is defined by two Splice Bubbles from *x*_1_ to *x*_5_. The primary and alternative paths of the bubbles are depicted by purple and orange coloring of the edges, respectively. (D) Comparison of ASE quantification with IsoTools from IsoSeq and with rMATS from Illumina RNA-Seq, for the untreated hepatocytes separately for the event classes: exon skipping (ES), intron retention (IR), mutually exclusive (ME) and 5’ and 3’ alternative splice sites combined (AS). (E) Differential splicing of SLC39A14 mutually exclusive exon 4 upon VPA treatment. Top track represents the reference annotation, and bottom tracks show log scaled sashimi plots of IsoSeq transcripts. Purple arcs represent splice junctions involved in the alternative splicing event, and are labeled with the total number and the fraction of supporting reads IsoSeq full length reads. Other splice junctions which are supported by at least 1% of the transcripts are depicted as green arcs, and by less than 1% grey.

In total, we identified and quantified 3.706 IR, 2.813 ES, 1.582, 3AS 1.319 5AS, and 369 ME events in the primary human hepatocytes, covered by at least 100 reads over both VPA and CTL, where the alternative contributed at least 10% of the reads. To validate the quantification of these events based on LRTS, we tried to match the LRTS IsoTools events with those identified by rMATs using the RNA-Seq data, but found only 41.6% of the events. In particular alternative events not covered by the reference annotation were not identified from the short reads: from the novel events, only 10.77% where also found by rMATS. In order to yield better overlap, we exported the events detected by IsoTools from LRTS to rMATS [16]. Since the rMATS event definition is less flexible, not all events could be exported, but 80.1% of the found IsoTools events could be exported to rMATS, and were quantified with the short reads. We found high correlation of PSI values for ES (r=.84), 5AS (r=0.8) 3AS (r=0.74) and IR (r=.62), while quantification of ME events were not correlated (r=-.05) (Figure 6 D). This result suggests that LRTS provides reliabe ASE detection and quantification, also for novel splicing. In contrast, short-read sequencing ASE detection preforms well if the splicing event is annotated, but is less reliable for novel events not present in the reference annotation.

### VPA induces differential splicing events in primary human hepatocytes on different levels of complexity

IsoTools provides a couple of statistical approaches to identify differential splicing events either with two samples or groups of samples. To demonstrate the capability of these approaches to identify biologically relevant events, we applied a two proportions z-test (cf. Methods) to compare VPA-treated hepatocytes to control samples, and found 806 differential splicing events in 556 different genes at an FDR of 1%. These events include 26 mutually exclusive exons, 259 exon skipping events, 105 5’ and 62 3’ alternative splice sites and 354 intron retention events. Also for the differential events, short read quantification with rMATS was in good agreement, with a Spearman correlation of ΔPSI of 0.68 (Supplementary Figure S6). In addition, we found differential usage of 974 transcription start sites and 306 poly-A sites between VPA and CTL. Differential splice events between VPA and CTL treated hepatocytes are listed in Suppl. Table 3.

These numbers suggest a widespread effect of VPA on the usage of TSS. Indeed, it has been shown that HDAC inhibitors (as well as DNMT inhibitors) specifically introduce cryptic transcription start sites (TSSs) in long terminal repeats [29]. This was shown for the HDAC inhibitor SAHA, a class I, II and IV inhibitor, while VPA is acting on class I and II proteins.

While mutual exclusive exon events are rare (3.2%), the most significant splicing event affects the mutually exclusive exons 4A and 4B of SLC39A14 (Solute Carrier Family 39 Member 14), a metal cation transporter responsible for the uptake of trace elements such as zink, iron and manganese in the liver [30]. Both versions, containing either exon 4A or 4B, yield functioning proteins, but the uptake kinetics vary substantially [31]. While we found both variants expressed at comparable level in the CTL sample, the proportion of 4B increased to 75% in the VPA treated sample (Figure 6E). A similar shift has been observed between normal and colorectal cancer samples (CRC) [32]. Notably, we found the same trend with rMATS using the short read RNA-Seq data when providing the events identified with IsoTools (42% to 70% PSI of 4B), while rMATS alone did not identify the correct event. The human liver samples where in-between the two hepatocyte samples (64% PSI of 4B).

Exon skipping events account for 32.1% of differential splicing. The most significant differential ES event affects increased inclusion of exon 32 of the FN1 gene. (fibronectin 1; *q* − *value* = 3, 55*E* − 111, PSI 9% in CTL and 15% in VPA). Again, rMATS analysis of short read RNA-Seq confirmed this differential event (PSI in CTL 12% and VPA 18%), while in human liver the exon was almost completely skipped (3% PSI). FN1 is one of the first genes for which alternative splicing was described, and regulation and functional effects of splicing in FN1 have been studied extensively [33]. The particular exon subject to the skipping event is referred to as extra domain A (EDA), one of two extra domains of this gene, and inclusion is regulated by Serine/arginine-rich splicing factors SRSF1 and SRSF3. While the extra domains are essential for normal development, elevated inclusion of EDA is associated with several diseases, including psoriasis, rheumatoid arthritis, diabetes and cancer. Previous studies observed similar effects on splicing of FN1 triggered by HDAC inhibitor sodium butyrate (NaB), but in this case resulting in elevated inclusion of exon 23, which is extra domain B[34]. Aberrant splicing of FN1 may be related to hepatotoxic effects, as downregulated or dysfunctional SRSF3, and subsequent aberrant splicing of its targets including elevated EDA-Fn1, has been associated with liver disease [35]. We thus conclude that the implemented LRTS workflow reliably identifies differential splicing events on different levels of complexity.

## Discussion

Here, we presented IsoTools, a flexible and powerful Python framework for the analysis of LRTS data. It provides data structures to search, access, and filter the transcripts, as well as functionality to compute quality control metrics, to compare and annotate the transcripts with reference annotations, to integrate data from several LRTS experiments, to quantify isoform expression levels based solely on LRTS, to perform statistical analysis for differential splicing, and to export of the data in several output formats. In addition, the tool facilitates the depiction of summary statistics as well as complex splicing models of individual genes and transcripts. IsoTools is explicitly designed to facilitate integration of custom metrics and algorithms to enable advanced users the application to novel use cases, such as for the analysis of single cell LRTS.

To characterize novel transcripts, we introduce a fine grade biologically motivated classification scheme, refining the established technically defined classes. The categories facilitate direct interpretation of differences between samples, and may hint towards specific disturbed splicing mechanisms, such as introduced by SF3B1 hotspot mutations, which specifically result in shifted in 3’ splice sites [36]. By far, the most common category of novel transcripts was “novel combinations of known splicing events”. Often, these combinations are separated by several kilobases, and thus cannot be identified with short read sequencing. Some of the identified novel transcripts result from misalignment of long reads at short exons. [37]. IsoTools helps identifying these cases, such that alignment algorithms can be improved in this regard.

Alternative splicing events (ASEs) have previously been defined as bubbles in splice graphs [15]. According to this definition, a “complete” ASE summarizes all variants that share common start and end splice sites, but no splice sites in between those. While the focus on splice sites sounds intuitive, this definition neglects the lack of splicing within a variant, with unintended consequences. For example, an alternative 5’ splice site event includes the flanking 5’ exon – and potentially additional splicing within this exon – but not the flanking 3’ exon, since the 3’ splice site is the same for the variants. This is incoherent, but also joins potentially distinct ASEs, unnecessarily increasing the complexity.

We propose a novel graph-based approach to identify and classify alternative splicing, based on bubbles in the segment graph, ensuring common exonic segments on both ends of the event without extending to common splice sites. In addition to be more coherent, our definition has two practical advantages over the definition on splice graphs: first, ASEs are subdivided in different classes, which can be analyzed independently. Second, while the original definition on splice graphs combines a variable number of alternatives to an event, our definition decomposes the event in two sets of transcripts. This binary perspective can be motivated biologically, as primary and alternative sets reflect the binary choices of an alternative splicing event (Figure 6 C) and has practical benefits, simplifying subsequent statistical analysis of ASEs.

Our definition of ASEs provides the basis for statistical tests for the detection of differential splicing events, either between two samples or between two groups of samples. This approach is fundamentally different from differential expression analysis on transcript level, for which the well established framework based on negative binomial generalized linear models (GLMs) [38, 39] can be applied, also with LRTS data [14]. However, identification of differential splicing events (DSE) has several advantages over differential transcript expression (DTE). While DTE may be a combination of differential regulation on gene level and differential splicing, DSE focusses on local splicing regulation, and is independent from gene expression levels. Further, DSE can distinguish several related or independent splicing events on the same gene. As the individual events can be classified, DSE facilitates categorization of differential splicing. Last, DSE aggregates statistical power from all transcript isoforms covering the event. Even transcripts affected by technical artefacts, such as RTTS and fragmentation, may still provide useful information for event level analysis. On isoform level, these artefacts would result in distinct transcript isoforms, and thus further increase the complexity and disturb the analysis if not filtered out. Hence, to interpret and validate the effects of differential splicing, DSE analysis yields more concise results.

Much like gene expression, alternative splicing is subject to biological variability within samples and groups of samples of the same condition. Statistical tests that compare individual samples, such as the two proportion z test and the likelihood ratio test with binomial model implemented here, neglect this variability, making the analysis prone to false positive results. Thus, the third implemented test, the likelihood ratio test with beta-binomial model, facilitates the comparison of groups of samples. The biological variability within the groups is estimated from the data and taken into account. While this approach promises to be more robust, it depends on replicates and thus could not be demonstrated with our data. LRTS datasets with biological replicates are the exception today, but continuously falling sequencing cost, higher throughput in combination with sample multiplexing, as well as better software facilitating additional applications will improve the cost benefit ratio of biological replicates.

In addition to DSE analysis, the identified alternative events can be used for exploratory analysis. By computing a PCA or UMAP embedding, the relation of samples with respect to event classes can be depicted. Furthermore, the events can be exported to be used with external tools designed for short read RNA-seq analysis. This facilitates integration and comparison of long and short read technologies.

We observed a depletion of short transcripts with the IsoSeq technology. The cause of this bias needs to be investigated. If it cannot be avoided technically, a dedicated normalization method would improve the accuracy of quantitative interpretation.

In primary human hepatocytes, IsoTools identified aberrant splicing events in different categories, caused by the HDAC inhibitor VPA. The role of HDACs in modulating alternative splicing has recently been emphasized by investigating the role of histone marks in the choice of splice sites and regulation of splicing [40]. We observed changes in splicing after VPA treatment on all levels of complexity but most prominently with usage of mutually exclusive exons, exon skipping and alternative TSS events, and we validated these events by short-read RNA-Seq of the same samples. While not yet reported after VPA treatment, many of the identified differential events have been observed to be triggered also by other HDAC inhibitors in comparable models, demonstrating the ability of LRTS to detect differential splicing between samples.

IsoTools is simple to install, flexible, versatile, and easy to use. It offers novel functionality, including expression quantification and differential splicing analysis, extending the range of potential applications for LRTS. Several example workflows cover relevant use cases and can simply be adapted by the user to realize more specific analysis. We demonstrated the utility of our tool by analyzing LRTS data from hepatocytes treated with VPA, and identified novel and differential splicing events, of which several are expressed also in human liver samples and are thus likely relevant *in-vivo*.

## Methods

### VPA-treated and control primary hepatocyte samples

The preparation of the samples processed here are described in [41]. In brief, human hepatocytes were exposed to 15 mM valproic acid (VPA) or 1% EtOH (CTL) and sampled after one day, two days and three days of treatment. For the 4th timepoint, VPA was washed out after day three, and cells were resampled after three additional days.

### PacBio IsoSeq Sequencing

For each time point and condition, we prepared triplicate cDNA samples. For library preparation, all VPA treated samples as well as the control samples were pooled. The libraries were sequenced on the PacBio Sequel II platform, using one 8M SMRT cells for each pool.

After confirming sample integrity using Agilent Bioanalyser, 300 ng RNA from each triplicate sample was incubated with NEBNext Single Cell RT Primer Mix for 5 min at 70 °C. Afterwards, Single Cell RT buffer and Single Cell RT enzyme Mix were added and incubated for 75 min at 42 °C. Then, template switching oligo was added, followed by 15 min of incubation at 42 °C. cDNA was cleaned up using 1:1 Pronex beads followed by two washing steps with 200 µl 80% ethanol and elution of the sample with EB. After addition of NEBNext Single Cell cDNA PCR master mix, NEBNext primer and the Iso-Seq Express cDNA PCR primer, cDNA was amplified in 12 cycles with 3 min elongation time. We targeted for standard transcript length of 2kb using 86 µl of Pronex beads for cleanup.

For sequencing, all control and VPA-treated samples were pooled equimolarly and PacBio IsoSeq libraries were prepared using the Express TPK 2.0 kit, including damage repair, end repair and A-tailing steps. Next, overhang adapters were ligated and the resulting libraries were cleaned up using 1:1 ProNex beads. Agilent bioanalyser assessment reported an average library insert sizes of 4,222 bp for CTL and 4,336 bp for VPA treated samples, respectively. Sequencing complexes were generated using Sequel II Binding Kit 2.0, Sequencing Primer v4.0, and Sequel II Polymerase 2.0, and then purified using ProNex beads.

Libraries for the pooled CTL and VPA treated samples were loaded individually together with PacBio internal control libraries on two Sequel II SMRT cells (diffusion loading). Sequencing was conducted with 30 h movie time, yielding 660.7 GB (CTL) and 460.1 GB (VPA) of total sequencing data. This resulted in 6,720,864 and 3,926,050 polymerase reads, with an average length of 98,305 and 117,202 bases.

### PacBio IsoSeq pre-processing

IsoSeq subreads were processed using the Iso-Seq v3.4 bioinformatic pipeline with recommended parameters. In brief, we used the `ccs` tool to call circular consensus sequences, lima to remove primers and adapters, and `isoseq refine` to filter out reads not featuring poly-A sequences. This resulted in 2,643,404 and 4,265,020 “full length non chimeric” (flnc) HiFi poly-A reads, with an average length of 3,636 and 3,880 bases for the VPA and CTL samples, respectively. 96.6% and 95.5% of these HiFi reads have an error rate of less than 1%, according to base quality values. The flnc reads were directly (without an additional clustering step) aligned to the human genome GRCh38.p13, obtained from the GENCODE website [42], by calling minimap2[43] with the `pbmm2 align --preset ISOSEQ` command. 2,611,571 and 4,196,197 of the reads could be aligned in one consecutive alignment, 30,561 and 66,293 were split in two or more parts, 1,272 and 2,530 were not aligned. All further analysis steps were performed with the IsoTools package, as described in the respective results sections. For these samples, the analysis including data import, computation of quality control metrics, characterization of novel isoforms, and differential analysis took about 67 minutes on a single CPU core, using a maximum of 20 GB RAM.

### Definition and classification of binary alternative splicing events

In analogy to the commonly used definition on splice graphs [15], we define binary alternative splicing events (ASEs) based on bubbles in the segment graph of a gene. In a segment graph, nodes represent disjoint exonic segments, and edges imply that the two exonic segments succeed one another in one or more transcripts. The segment graph is bi-directed, as each node has a set of incoming and outgoing edges, and ordered by the genomic position, meaning an edge from node *x*_*i*_ to node *x*_*j*_ is only allowed if *x*_*i*_ < *x*_*j*_, e.g. the genomic end position of *x*_*i*_ is smaller or equal to the start position of *x*_*j*_. If two succeeding segments are separated by an intron, the edge represents a splice junction. On the other hand, if the genomic end position of the preceding segment corresponds to the genomic start position of the succeeding segment, the edge is called internal. This implies either an alternative preceding or succeeding segment, connected by a splice junction edge.

Bubbles are structures in the segment graph, with two paths starting in a common segment *x*_*s*_ and ending in a common segment *x*_*e*_, but the paths do not share any segments except *x*_*s*_ and *x*_*e*_ (cf. Figure 6A-C). We define the “primary path” as the path for which the outgoing edge from *x*_*s*_ exceeds the outgoing edge of the other path, which in turn is called the “alternative path”. We further categorize the alternative path in 5 different classes: the alternative is classified as mutually exclusive (ME) if the primary path from *x*_*s*_ to *x*_*e*_ contains at least one additional segment *x*_*ME*_. The definition of the other classes depend on whether the outgoing edge of *x*_*s*_ and the incoming edge of *x*_*e*_ on the alternative path correspond to splice junctions or internal edges (within an exon). Alternative paths are called exon skipping (ES) if both edges are splice junctions, and as intron retention (IR) if both edges are internal. If one of the edges is a splice junction and the other internal, the alternative is classified as 5’ or 3’ alternative splice site (5AS and 3AS). For each primary path, we group all alternative paths of the same category, and find the set of transcripts *A* supporting one of the alternatives and *B* supporting the primary.

This definition results in a finite set of classified binary alternative splicing events for each gene. They can be quantified by the proportion spliced in (PSI), defined as the number of long reads supporting transcripts from *A* over the total number of reads, supporting transcripts from *A* or *B*.

### Statistical tests for differential splicing

Detecting differential splicing events between two samples or groups of samples is a major goal of splicing analysis. This can be achieved by specific statistical tests. In IsoTools, we implemented 3 statistical tests for differential splicing. The first two, two-proportions z-test and likelihood ratio test with binomial model, apply if two individual samples are compared. For the third test, splicing proportions are modeled with beta-binomial distributions. With this approach, the likelihood ratio test is appropriate for the comparison of two groups of samples, as the models account for variability within groups.

The **two-proportions z-test** is specified by the statistic

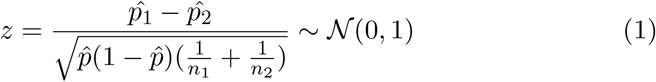

where *n*_*i*_ = *k*_*i*_ + *l*_*i*_ is the total number of reads of sample *i* ∈ [1, 2] covering the event. *k*_*i*_ and *l*_*i*_ are the number of reads supporting the alternative (set *A*) and primary (set *B*) variants, respectively. 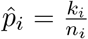 is the proportion of reads supporting the alternative for sample *i*, and 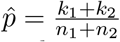 the proportion supporting the alternative for both samples combined.

Alternatively, when the number of supporting reads is modeled with a binomial distribution, a **binomial likelihood ratio test** can be applied.

This test is specified by the statistic

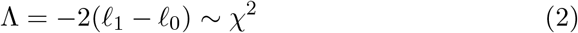

where

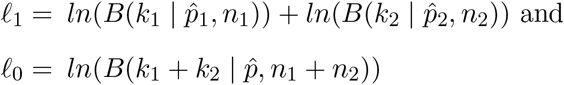

are the maximized log-likelihoods under the alternative *H*_1_ and the null hypothesis *H*_0_. *B*(*n* | *p, n*) is the probability mass function of the binomial distribution, which is maximal at the empirical proportion 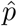 defined above. Both tests yield very similar results (p-values r=0.999, see Suppl. Figure S7).

In the presence of two groups of samples IsoTools offers a **beta-binomial mixture likelihood ratio test**. This test models the variability within the two groups with a beta-binomial mixture distribution, a binomial distribution where the probability parameter *p* follows a beta distribution *Beta*(*a, b*). The maximum log-likelihood parameters *â* and 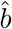 are determined numerically by a quasi Newton optimization method (LM-BFGS from SciPy [44]).

In addition to ASEs, all tests defined above can also be applied to detect differential usage of transcription start and poly-A sites. In this case, all transcripts supporting a particular start/poly-A site are considered the alternative set *B*, whereas all other transcripts constitute the primary set *A*.

### RNA-Seq

RNA-Seq data for the primary hepatocytes and liver samples were downloaded from ENA accession PRJEB22198 and PRJEB35350 respectively. The short reads were aligned to the human reference genome GRCH38.p13 using `STAR` aligner version 2.7.6a [45], with provided gff annotation from GENCODE release 36 including annotation of non-chromosomal scaffolds [42]. For hepatocytes, we merged all alignments from the same time point, to resemble the IsoSeq samples. We used `rsem` v1.3.1 [46] with the transcriptome alignment from STAR to obtain read counts on transcript and isoform level. Next, we used rMATS `rmats-turbo` v4.1.1 [16] to find differentially spliced events between VPA-treated hepatocytes and control, either using events calculated by rMATS, or by providing events generated by IsoTools from the GENCODE reference as well as from IsoSeq LRTS.

### ENCODE CAGE data

We downloaded CAGE TSS peaks for HepG2 cells from the ENCODE data portal [22] (https://www.encodeproject.org/) with the following identifiers: ENCFF089AFK, ENCFF220OWX, ENCFF241CGD, ENCFF248QKX, ENCFF373BNI, ENCFF419FNU, ENCFF875ILB, ENCFF885VJU.

## Data Availability

The primary human hepatocyte LRTS data is available from ENA under accession number PRJEB46194.

## Code Availability

IsoTools is available at https://github.com/MatthiasLienhard/isotools and the PyPI software repository. The documentation, including tutorials with example applications and case studies as well as the API reference, is available at https://isotools.readthedocs.io/en/latest/. The github repository includes a Jupyter notebook to replicate all analyses from this study.

## Acknowledgements

The study was financed by the German Research Foundation (DFG) with the grant HE4607/7-1 and the Federal Ministry of Education and Research with the grant SafetyNet (161L0242A).

## Notes

https://github.com/MatthiasLienhard/isotools

https://isotools.readthedocs.io/en/latest/

https://pypi.org/project/isotools/

### Competing Interest Statement

The authors have declared no competing interest.

