## Supplementary Figures for "Long-read transcriptome sequencing analysis with IsoTools"

### Long-read transcriptome sequencing analysis with IsoTools
### Supplementary Figures

Matthias Lienhard[1,*], Twan van den Beucken[2], Bernd Timmermann[3], Myriam Hochradel[3], Stefan Boerno[3], Florian Caiment[2], Martin Vingron[1], and Ralf Herwig[1,*]

[1] *Max Planck Institute for Molecular Genetics, Department of Computational Biology, Ihnestraße 63-73*

[2] *Maastricht University, Department of Toxicogenomics, Universiteitssingel 40, 6229ER Maastricht, NETHERLANDS*

[3] *Max Planck Institute for Molecular Genetics, Sequencing core unit, Ihnestraße 63-73, 14195 Berlin, GERMANY*

[*] *To whom correspondence should be addressed.*

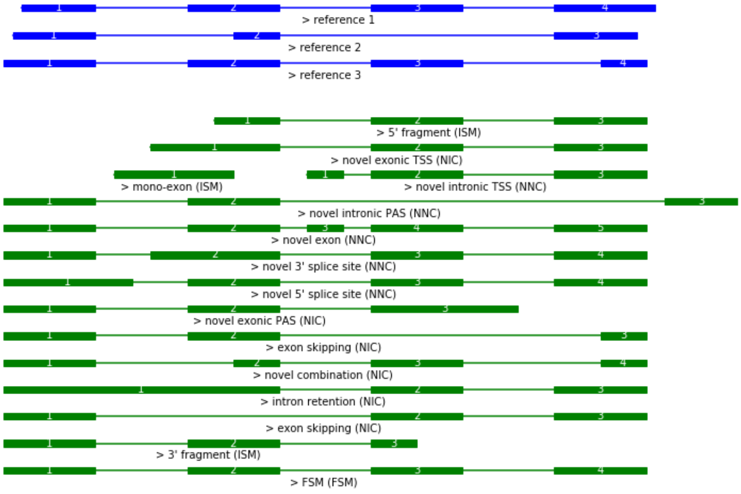


Figure S1: Prototypical examples for novel transcript classes. Reference transcripts are depicted in blue, and classified novel transcripts in green. Labels indicate the novelty class, with the Sqanti classification in brackets.

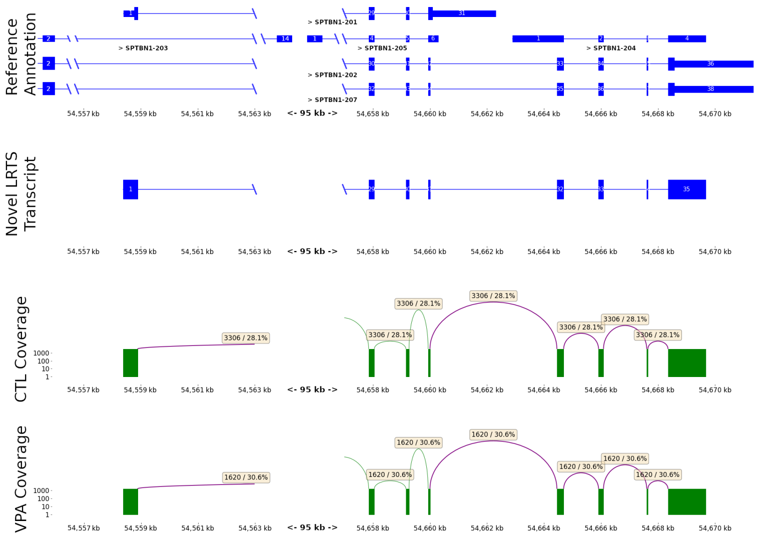


Figure S2: The novel isoform of the SPTBN1 gene is a combination from the TSS (left) of isoform SPTBN1-201 (first line in the top track) and the polyA site (right) of isoforms SPTBN1-202 and SPTBN1-207 (third and forth lines in the top tracks). The top gene track depicts the reference isoforms, and below the novel IsoSeq isoform. Coding regions in the reference transcripts are represented by broader blocks. The Sashimi plots in the bottom represent the coverage of this isoform for CTL and VPA hepatocyte samples respectively.

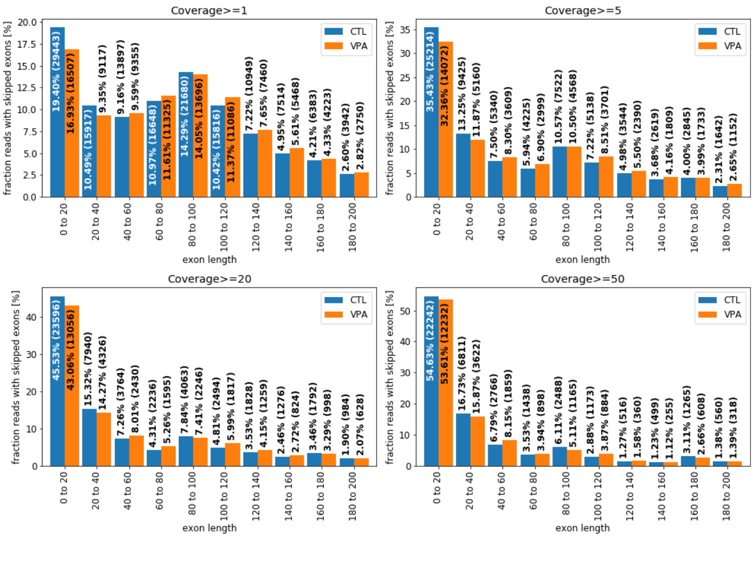


Figure S3: Histogram of novel exon skipping transcripts for different ranges of exon lengths. Short exons are not correctly aligned by the alignment tool, resulting in the detection of false skipped exons.

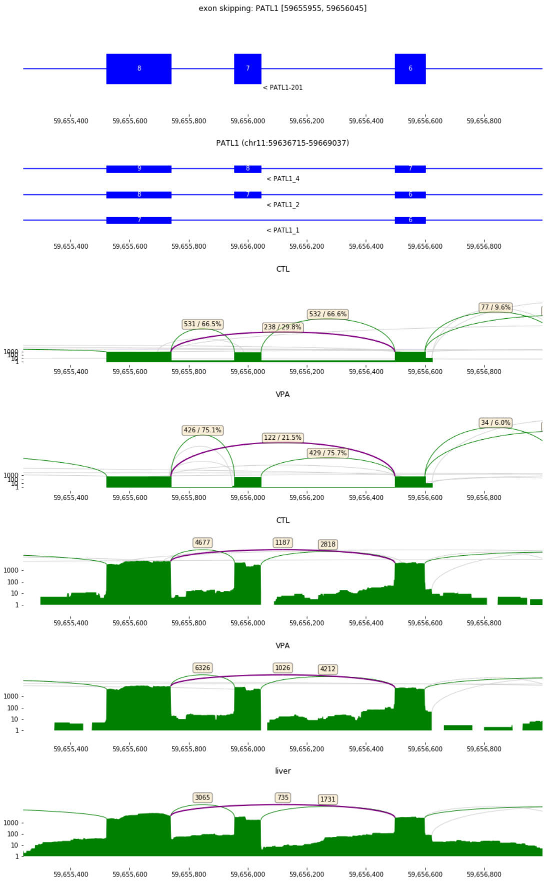


Figure S4: The novel isoform of PATL1 skipps exon 7 of the GENCODE annotation (top track). Relevant isoforms detected by IsoSeq are depicted in the second track. The following two Sashimi plots depict the IsoSeq coverage for the hepatocytes (CTL and VPA), and the bottom three Sashimi plots Illumina RNA-Seq coverage. The exon skipping junction is depicted in purple, other highly covered junctions in green, and poorly covered junctions in grey.

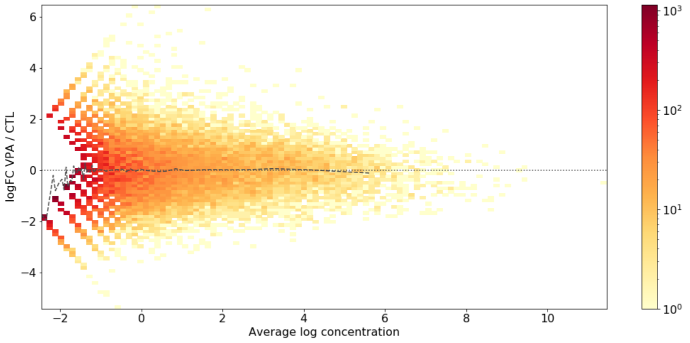


Figure S5: MA plot of upper quantile normalized IsoSeq transcript read counts in VPA vs CTL treated hepatocytes. Dashed line represent the binned average of logFCs, within equally sized bins of 500 transcripts.

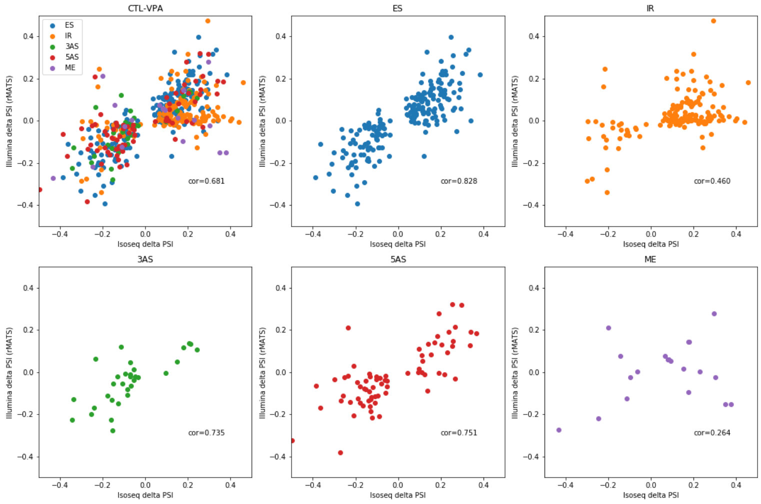


Figure S6: Comparison PSI differences between VPA treated and control samples for differential ASE (two-proportions z-test $FDR < .01$), quantified by IsoTools from LRTS and rMATS from short read RNAseq, for all events and the splicing event classes individually.

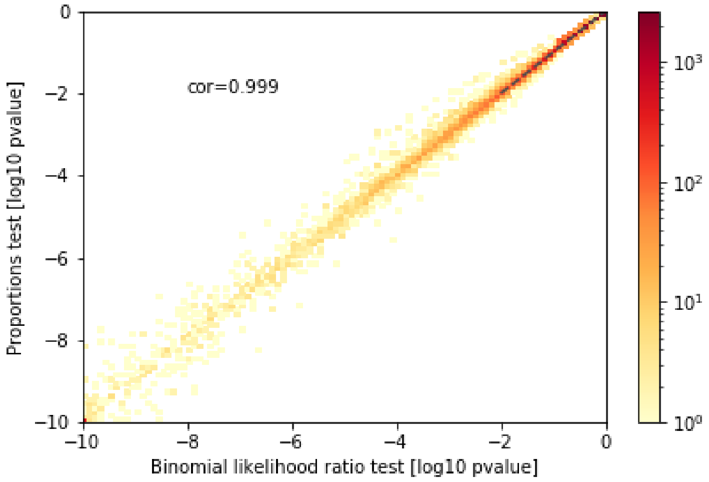


Figure S7: Comparison of binomial likelihood ratio test and two proportions z-test, for VPA vs. CTL samples.
